# Construction of a randomly barcoded insertional mutant library in the filamentous fungus *Trichoderma atroviride*

**DOI:** 10.1101/2025.11.30.691285

**Authors:** Lori B. Huberman, José M. Villalobos-Escobedo, Jeffrey M. Skerker, Ran Shi, Adriana M. Rico-Ramírez, Catharine Adams, Adam P. Arkin, Adam M. Deutschbauer, N. Louise Glass

## Abstract

Filamentous fungi play key roles in ecosystems, agriculture, biotechnology, symbiosis, and disease, yet the large-scale characterization of gene function in these organisms remains limited by low transformation efficiencies and their multinucleate, syncytial cells, which complicate high-throughput screening strategies. To address the challenge of high-throughput screening in filamentous fungi, we developed methods to construct a genome-wide barcoded insertional mutant library in *Trichoderma atroviride*, a filamentous fungus widely used as a biocontrol agent against bacterial and fungal plant pathogens. Our strategy leveraged randomly barcoded transfer DNA insertions from plasmid libraries containing hundreds of millions of unique DNA barcodes and a broad host-range drug resistance marker delivered via *Agrobacterium tumefaciens* into *T. atroviride*. By optimizing transformation conditions, we achieved up to 600 independent transformants per infection event, resulting in a library of over 31,000 mapped insertions disrupting 7,104 of the 11,863 predicted genes in the *T. atroviride* genome. This resource establishes a scalable platform for high-throughput functional genomics in filamentous fungi, enabling both fundamental investigations of fungal biology and engineering approaches toward improved medical applications, biotechnology, and sustainable agriculture.

## Introduction

Filamentous fungi are among the most versatile organisms on Earth, with crucial roles in ecosystem dynamics, agriculture, biotechnology, and human health. They degrade complex carbon sources, including lignocellulosic biomass and plastics, contributing to carbon cycling in the environment and bioremediation (1–3). They also produce a vast array of secondary metabolites and pigments (4) and are capable of remarkable developmental transitions in response to environmental cues, including light and pH (5–8). Filamentous fungi are also plant, human, and animal pathogens responsible for global crop losses (9) and threats to human and animal health (10, 11). Despite their biotechnological, medical, agricultural, and ecological relevance, our ability to characterize gene function in filamentous fungi using high throughput methods remains limited.

Randomly barcoded transposon or transfer DNA (TDNA) insertional mutagenesis has enabled the systematic discovery of gene function in bacteria and unicellular fungi, including *Rhodosporidium toruloides* (12–15). This method uses transposons or TDNA elements carrying unique DNA barcodes that are randomly inserted across the genome of the organism under study, thus creating an insertional mutant library. Exposing this mutant library to experimental conditions of interest enables evaluation of the fitness of thousands of mutants in parallel using PCR and high-throughput sequencing by measuring changes in the relative barcode abundance across experimental conditions. This technique has significantly accelerated gene annotation and systems-level understanding in microbial models, including during host-microbe interactions (16).

Although high-throughput functional genomics strategies have been successfully applied to uninucleate microbes, it remains a significant challenge in filamentous fungi. The complexity of multinucleate mycelial growth in these organisms, combined with low transformation efficiencies and limited genetic tools, has hindered the development of high-throughput functional genomics approaches (17, 18). To develop these tools in filamentous fungi, we developed a high-efficiency randomly barcoded insertional mutagenesis sequencing pipeline (RB-TDNAseq) and applied it to the soil fungus *Trichoderma atroviride* IMI 206040, a well-established biocontrol agent that promotes plant growth and suppresses phytopathogenic fungi (19, 20). We first constructed large-scale plasmid libraries with hundreds of millions of unique barcodes and a fungal-selectable marker optimized for *Agrobacterium tumefaciens*-mediated transformation. We subsequently optimized *A. tumefaciens*-mediated transformation of *T. atroviride*, thereby generating a library of 31,422 mapped insertions, resulting in disruptions of 7,104 predicted genes in the *T. atroviride* genome. Our results demonstrate a scalable technique for genome-wide functional analysis in filamentous fungi applicable to a wide range of species, opening new avenues for fundamental research and applied biotechnology in these multicellular and multinucleate organisms.

## Results

### Developing protocols to construct genome-wide insertional mutagenesis libraries

Deploying RB-TDNAseq in filamentous fungi has several challenges relative to most unicellular microbes, including low transformation efficiencies, relatively large genomes (∼40-150 Mb in a typical filamentous fungal genome compared to ∼12 Mb for *Saccharomyces cerevisiae* or 0.1-10 Mb in bacteria), increased gene content, larger intergenic regions, and multinucleate cells. To reach our goal of a library of mutants with uniquely barcoded insertions in every nonessential gene in a filamentous fungal genome, we first created a library of hundreds of millions of plasmids containing a drug resistance cassette each tagged with a unique DNA barcode (Fig. 1) in a plasmid backbone optimized for *A. tumefaciens*-mediated transformation (21). As a selectable marker, we chose to use a hygromycin resistance cassette (22) suitable for transformation in a variety of filamentous fungi (23). We first optimized an established Golden Gate Cloning protocol (24) for high efficiency cloning, enabling the generation of a library of an estimated 1 billion uniquely barcoded plasmids in *Escherichia coli* (Dataset S1). We subsequently optimized plasmid transformation of a strain of *A. tumefaciens* resulting in a library containing 140 million uniquely barcoded plasmids with a hygromycin resistance marker for transformation of a wide variety of species of filamentous fungi (Dataset S1) (see Materials and Methods).

**Figure 1.**
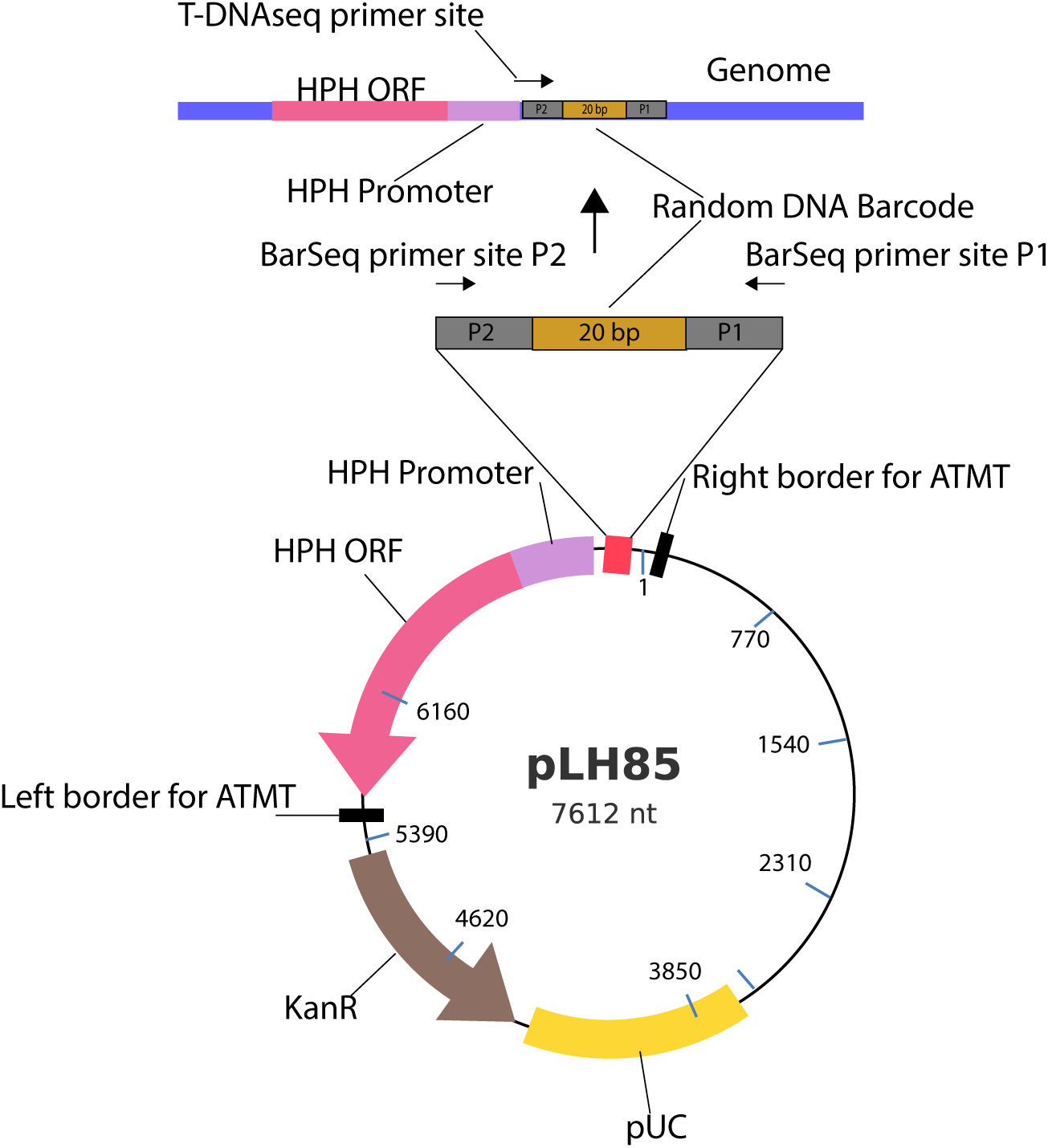
Circular map of the pLH85 plasmid containing the hygromycin B resistance cassette (*HPH ORF*). The plasmid has a total size of 7,612 bp. The *pUC* region is highlighted in yellow, the kanamycin resistance gene for selection in *Escherichia coli* and *Agrobacterium tumefaciens* in brown, the *hph* gene for selection in fungi in pink, and the *hph* gene promoter in purple. The borders for ATMT indicate the region of the plasmid that is the TDNA insertion into the fungal genome, which includes the hygromycin B resistance cassette and the 20 bp random barcode. The TDNAseq primer site for sequencing the insertion sites in the *T. atroviride* genome and the barcode sequencing primers for future fitness assays are indicated. Primer sequences are listed in Table S1.

Generating insertional mutagenesis libraries in filamentous fungi with tens to hundreds of thousands of unique insertions requires a high-efficiency transformation protocol that can be performed in a high-throughput manner. *A. tumefaciens*-mediated transformation (ATMT) is possible in a wide range of fungal species (25) and is more amenable to high-throughput implementation than other protocols widely used across fungal species (i.e., protoplast transformation) (26). We optimized ATMT of *T. atroviride* asexual spores (conidia) by altering the concentration and pH of the *A. tumefaciens* induction media buffer, the concentration of the metabolite (acetosyringone) that induces TDNA transfer by *A. tumefaciens*, the age of the *T. atroviride* conidia, the length of co-incubation between *A. tumefaciens* and *T. atroviride*, and the temperature at which the co-incubation was performed (See Materials and Methods; https://www.protocols.io/view/transformation-protocol-atmt-trichoderma-atrovirid-261gekk87g47/v1).

Adjusting these variables allowed us to increase our transformation efficiencies to approximately 600 transformants per transformation event.

Conidia of many *Trichoderma* species are uninucleate (27, 28), thus enabling the recovery of homokaryotic transformants following ATMT. However, even with the addition of selective antibiotics, bacterial DNA was present in initial library preparations. To eliminate contamination by *A. tumefaciens* DNA, we forced *T. atroviride* transformants to invasively grow through selective top agar prior to collecting homokaryotic conidia from transformants (Materials and Methods; https://www.protocols.io/view/transformation-protocol-atmt-trichoderma-atrovirid-261gekk87g47/v1)*. A. tumefaciens* cells were trapped under the top agar, resulting in clean separation of *T. atroviride* transformants from *A. tumefaciens* cells. A schematic of the transformation protocol is shown in Fig. 2.

**Figure 2.**
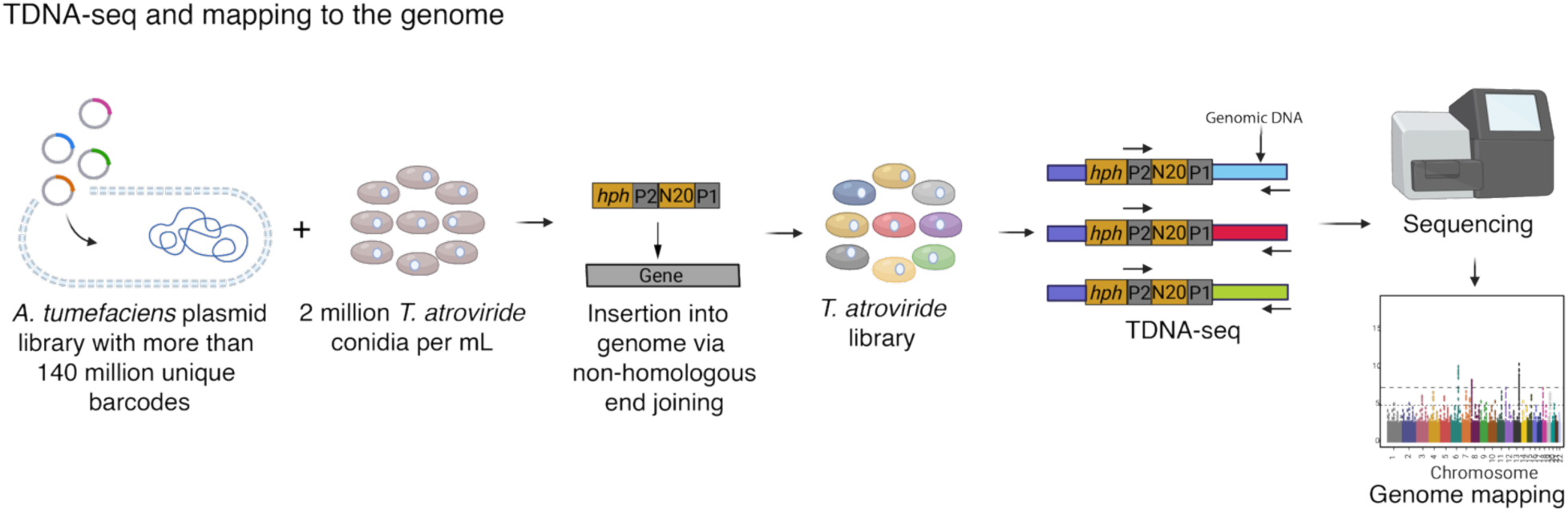
Schematic representation of RB-TDNAseq in *T. atroviride*. Generation of a barcoded insertional *T. atroviride* mutant library using *A. tumefaciens*-mediated transformation. Plasmids containing fungal drug resistance (*hph*) cassettes, which confers hygromycin resistance, tagged with an estimated 1 billion unique DNA barcodes (N20), flanked by common PCR priming sites (P1 and P2), were generated using Golden Gate Cloning and introduced into *E. coli*. These plasmids were then extracted from *E. coli* and transformed into *A. tumefaciens*, creating a library of approximately 140 million uniquely barcoded plasmids for *T. atroviride* transformation. Following selection of transformed *T. atroviride* homokaryotic transformants via hygromycin resistance, genomic DNA was extracted, and insertion sites were sequenced to map the barcodes to their respective genomic locations (blue, red, and green lines indicate the genomic region sequenced to determine the TDNA insertion site).

### The distribution of insertions is random across the *T. atroviride* genome

Following ATMT, hygromycin-resistant *T. atroviride* transformants were inoculated onto potato dextrose agar (PDA) plates supplemented with a number of amino acids, nucleic acids, and vitamins (alanine, arginine, asparagine, aspartic acid, cysteine, glutamic acid, glutamine, glycine, histidine, isoleucine, leucine, lysine, methionine, phenylalanine, proline, serine, threonine, tryptophan, tyrosine, valine, p-aminobenzoic acid, adenine, inositol, and uracil: PDA+) and allowed to sporulate. Conidia were subsequently collected for DNA extraction. A modified version of the KAPA DNA sequencing library preparation protocol for Illumina sequencing was used, in which the PCR amplification step was optimized to highly enrich for DNA fragments containing a barcoded insertion (29) (Materials and Methods). After adapter ligation to fragmented genomic DNA, we performed two PCR enrichment steps for genomic DNA fragments containing a barcode with a forward primer immediately upstream of the barcode within the TDNA insertion cassette and a reverse primer in the Illumina adapter that was ligated to the *T. atroviride* genomic DNA downstream of the insertion site (Fig. 1). The randomly barcoded TDNA (RB-TDNA) libraries were subsequently sequenced on an Illumina MiSeq, allowing the association of barcodes with their insertion locations across the *T. atroviride* genome (Fig. 3).

**Figure 3.**
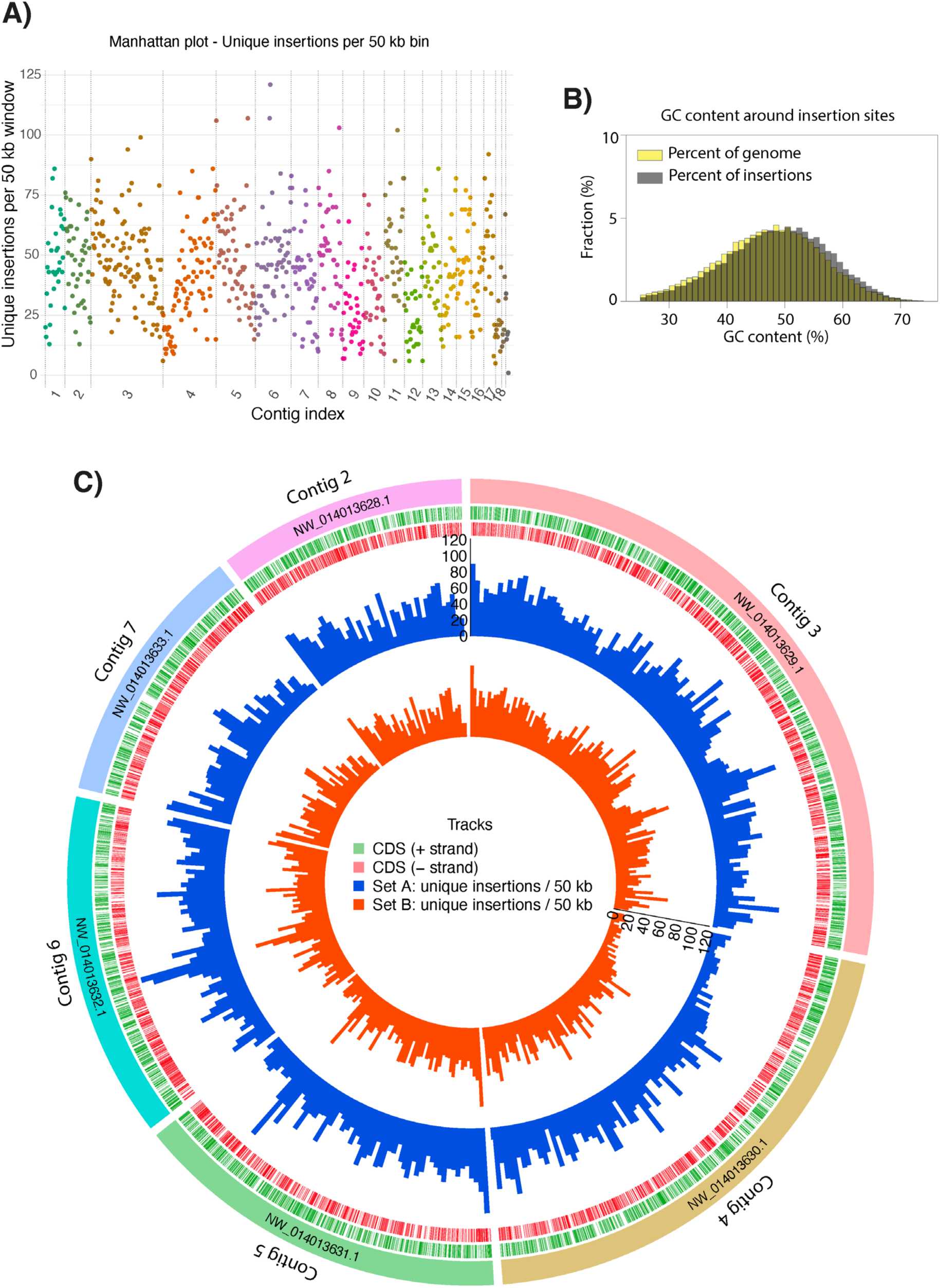
Distribution of TDNA insertions occurs randomly across the *T. atrovirid*e genome. (**A)** Manhattan plot illustrates the number insertions containing unique barcodes that mapped across the 20 largest *T. atroviride* contigs (https://mycocosm.jgi.doe.gov/Triat2/Triat2.home.html). (**B**) Histogram of the percent of insertions in regions of the genome with a particular GC content (grey) compared to the percent of the genome with a particular GC content (yellow). The GC content of an insertion site is defined as the 100 bp region surrounding the insertion site. (**C**) Circos plot showing the genomic distribution and density of uniquely barcoded insertions identified in two independent KAPA DNA preparation and sequencing (KAPA Biosystems) runs on the TDNA insertional mutant library (Set A, blue; Set B, orange). Each bar represents the number of unique TDNA insertion sites within non-overlapping 50 kb genomic windows. The y-axis corresponds to the absolute count of unique insertions per window, and both libraries share the same scale for direct visual comparison. Gene coding sequences (CDS) are indicated in green for the positive strand and red for the negative strand.

The RB-TDNA insertion site sequencing data showed that 72.7% of the reads mapped to insertion sites in the *T. atroviride* genome with 16.2% of reads mapping to the TDNA plasmid (indicative of bacterial contamination, incorrect processing of the plasmid during ATMT, or concatemers formed through back-to-back TDNA insertions). The remaining 11.1% of reads did not contain barcodes (Fig. S1). In total, 31,422 unique barcoded insertions were mapped in the *T. atroviride* insertional mutagenesis library. These insertions exhibited a similar density across all contigs of the *T. atroviride* genome (30) (https://mycocosm.jgi.doe.gov/Triat2/Triat2.home.html) (Fig. 3A) and lacked detectable insertion bias due to GC content in a 100 bp window surrounding the insertion site (Fig. 3B). Data from two independent KAPA DNA sequencing preparations of genomic DNA harvested from the mutant library (set A in blue and set B in orange; Fig. 3C) also showed high similarity.

To examine the distribution of insertion sites in greater detail, we plotted the insertion profiles for the largest 20 contigs, which were each at least 200 kb long (Fig. S2). Insertions were distributed relatively uniformly across the contigs, as shown for the six largest contigs (Fig. 3C), except for a few hotspots. These insertion hotspots may denote unidentified genomic features that modulated accessibility of TDNA to chromosomal DNA (i.e., chromatin state or DNA-protein interaction). There were also regions within the contigs that showed low insertion density (Fig. 3C; Fig. S2).

### Analysis of RB-TDNA insertions in the mutant library

Although we did not detect an insertion bias at the genomic level, it is possible a bias in insertion events was associated with particular genomic features. To assess this possibility, we used a generalized linear model (GLM) with a gamma distribution to evaluate differences in the relative fraction of insertion locations in the library across five genomic features (terminator, promoter, intron, exon, and intergenic regions) compared to the relative fraction of those genomic features in the genome. The results showed significant differences in all categories (p < 0.05). The library had more insertions in predicted promoter sequences and intergenic regions and fewer insertions in predicted terminator regions, introns, and exons than would be expected by chance (Fig. 4A; Dataset S2).

**Figure 4.**
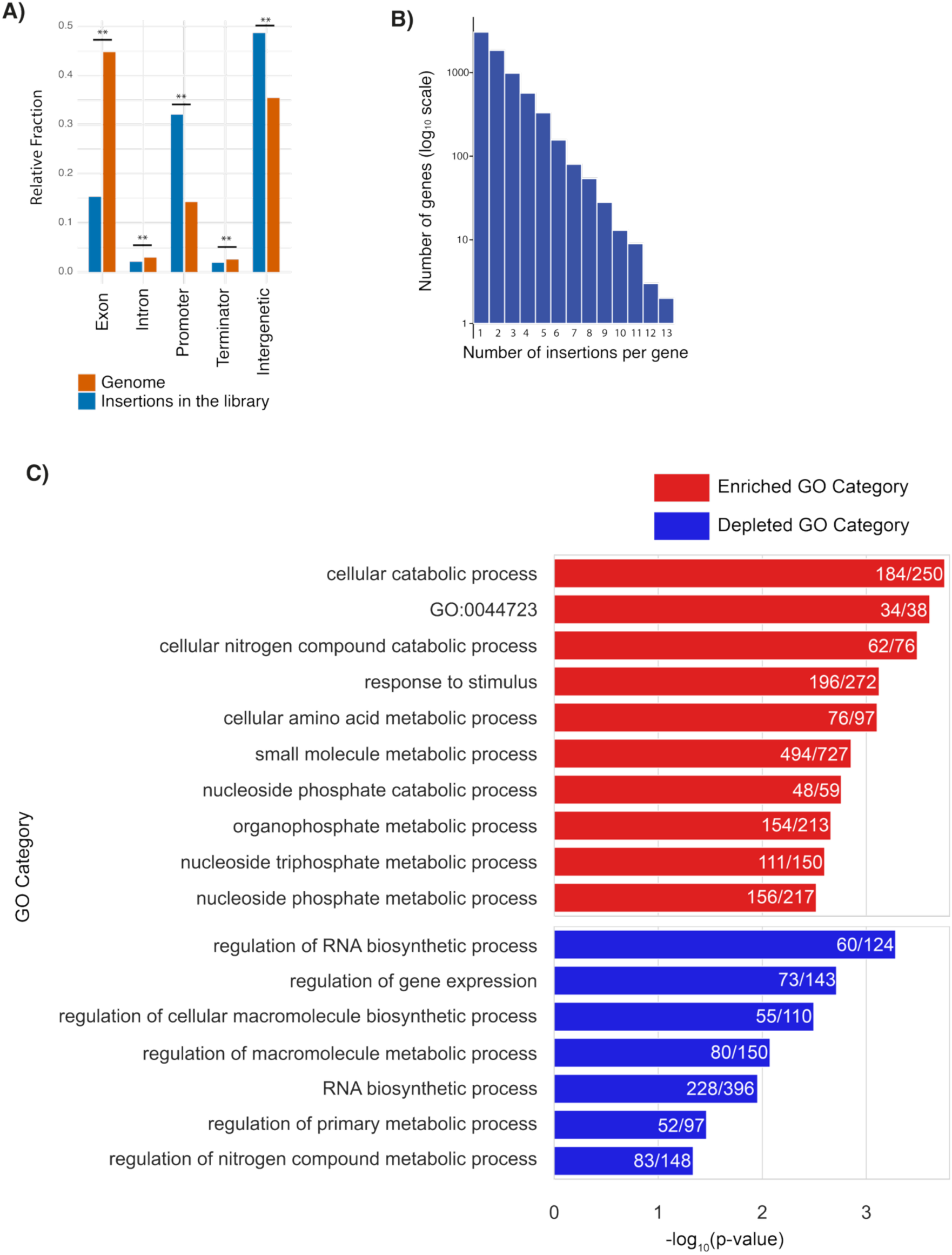
Insertions in the *T. atroviride* mutant library are biased towards promoters and intergenic regions. (**A**) Statistical analysis of the fraction of insertions found in exons, introns, promoters, terminators, and intergenic regions relative to the fraction of the genome in that genetic feature. A generalized linear model (GLM) with a gamma distribution was applied to identify significant differences (**p < 0.001). (**B**) Histogram of the number of insertions per gene. The y-axis shows the number of genes with a given number of insertions on a logarithmic scale (base 10), and the x-axis shows the total number of insertions in each gene for all genes with 13 or fewer insertions. Three genes with more than 13 insertions (14, 16, and 19 insertions) are not represented in this plot. (**C**) GO categories that are enriched and depleted in the insertion library. Fractions denoted on the bars represents the number of genes with insertions versus the total number of genes in the genome in that category, with bars indicating the p-values. Categories in red are enriched; categories in blue are depleted. The enriched categories with the 10 lowest p-values and all depleted categories with p-values < 0.05 are shown in the graph. For a full list of GO categories, see Dataset S3.

An analysis of predicted *T. atroviride* genes containing insertions revealed that 7,104 genes have at least one insertion, accounting for 60% of the 11,863 predicted genes in the *T. atroviride* genome (Fig. 4B). Thirty genes have 10 or more insertions, while 3,041 have only a single insertion (Fig. 4B). The gene with the largest number of insertions was an 8,033 bp gene predicted to encode a DNA-binding protein (ID: 283087; 19 insertions). A number of other genes that encode proteins predicted to bind DNA also had 10 or more insertions: a Rad50 homolog (ID: 252710; 13 insertions), a gene encoding a curved DNA-binding protein (ID: 298169; 10 insertions), three genes predicted to encode transcription factors (ID: 213474, ID: 322845, ID: 211488; 11 insertions each). In addition, several genes lacking annotation also contained a high number of insertions (including ID: 40706 and ID: 84521, with 16 and 13 insertions, respectively).

Functional annotation using Gene Ontology and enrichment analysis (GO; https://geneontology.org/docs/go-enrichment-analysis/) revealed that all biological process categories identified in the genome were represented by genes with insertions, with no evident bias toward any specific functional class (Fig. 4C; Dataset S3). However, we found that 33 categories, including *cellular catabolic process* (GO:0044248), *response to stimulus* (GO:0050896), and *cellular amino acid metabolic process* (GO:0006520), were significantly enriched for insertions within genes (FDR < 0.05). In contrast, seven functional categories appeared to be underrepresented in the library (fewer genes with insertions than expected by chance), specifically *regulation of RNA biosynthetic process* (GO:2001141), *regulation of gene expression* (GO:0010468), *regulation of cellular macromolecule biosynthetic process* (GO:2000112), *regulation of macromolecule metabolic process* (GO:0060255), *RNA biosynthetic process* (GO:0032774), *regulation of primary metabolic process* (GO:0080090), and *regulation of nitrogen compound metabolic process* (GO:0051171) (Fig 4C; Dataset S3).

## Discussion

The rapid increase in sequenced filamentous fungal genomes highlights the challenges of accurate functional gene annotation (31, 32). This issue stems largely from the difficulties in generating comprehensive, genome-wide datasets in filamentous fungi. Classical forward genetic screens of randomly generated mutants or reverse genetic screens of filamentous fungal deletion mutant collections (33, 34) are critical but are time-consuming tools to construct for functional gene characterization.

Here, we describe methods to establish RB-TDNAseq for high-throughput functional genomics in the filamentous fungus *T. atroviride*. In previous studies, ATMT and insertional mutagenesis has been used to identify genes in a few filamentous fungi including the plant pathogen *Magnaporthe orzyzae,* and *Trichoderma reesei* (35, 36), in addition to a variety of yeast species (14, 37, 38). However, previous applications of insertional mutagenesis in filamentous fungi have typically used a tailed-PCR approach to identify mutants, which is laborious when only used for a single screen and requires substantial sequencing depth (36). Here, we generated a randomly barcoded insertional library, which greatly expedites screening a mutant library under a variety of conditions. Our development of an *A. tumefaciens* plasmid library containing 140 million uniquely barcoded plasmids that can be used for transformation in a variety of fungi and containing a resistance marker (hygromycin B) that is widely used in filamentous fungi will greatly expedite functional analyses of filamentous fungal genomes, including comparative analyses. In this study, we highlight variables in the ATMT protocol important for increasing transformation efficiencies in filamentous fungi, facilitating the development of both insertional mutant libraries and improved targeted transformation protocols.

RB-TDNAseq libraries enable inexpensive high-throughput screens (12). After an initial deep sequencing step to map insertion locations and association of insertions with a unique DNA barcode, pooled fitness screens can be performed by PCR using universal primers that flank the barcode followed by DNA sequencing. Barcode sequencing requires lower sequencing depth than is typically necessary in an unbarcoded insertional mutant library and is therefore relatively inexpensive. However, performing pooled fitness screens in filamentous fungi has challenges when compared to uninucleate microbes, including yeast species (14, 37, 38), because of the filamentous fungal multinucleate syncytial mycelial growth habit, and the potential for cell fusion between both germinated asexual spores and between hyphae. In addition, because some species of filamentous fungi have multinucleate asexual spores, a purification step for the recovery of homokaryotic mutants must be performed. If spores are multinucleate, it is possible that complementation of a loss-of-function mutation caused by insertional mutagenesis could occur in heterokaryotic cells. The use of *T. atroviride*, which has uninucleate asexual spores, circumvented this issue. Strategies to harvest large numbers of uninucleate or homokaryotic transformants as well as methods to reduce the frequency of asexual fusion between hyphae (and thus complementation) during fitness experiments are an important consideration when applying this technology to other fungal species.

In this study, we focused our analysis on insertions in coding regions. However, our library contained more insertions in intergenic than in coding regions. Although we do not know the role that most noncoding DNA plays in filamentous fungi, many long non-coding RNAs (lncRNAs) and microRNAs have been identified in filamentous fungal genomes and several have known functions (39, 40). For example, in *Fusarium graminearum,* non-coding regions play a role in sexual reproduction (41), non-coding regions are involved in cellulase production in *T. reesei* (42), and microRNA-like RNAs play a role in an injury response in *T. atroviride* (43). It will be the role of future analyses to determine whether it is possible to use barcoded fitness assays to functionally characterize non-coding regions, including lncRNAs and microRNAs. Additionally, a recent study using transposon mutagenesis in *Cryptococcus neoformans* demonstrated that it is possible to assay functions of essential genes using insertions in promoter regions (38). Future studies could use this technique to investigate the roles of essential genes in filamentous fungi using this analysis technique.

We demonstrate a method and provide crucial tools for relatively inexpensive generation of a mutant library containing tens of thousands of mutants for pooled, high-throughput fitness assays applicable to filamentous fungi across the fungal kingdom. The barcoded mutant library we generated in this study paves the way for studying fungal-host interactions, responses to environmental stimuli, and interactions between fungi and other microbes. Recent studies suggest that these inter-kingdom interactions are far more interdependent and mechanistically intricate than previously thought. For example, fitness assays in bacterial species have revealed mechanisms of fungal-bacterial interactions in fermented cheese rinds (44) and upon exposure of bacteria to fungal exo-metabolites (45). Fitness assays of a targeted deletion collection of predicted protein kinases in *A. fumigatus* identified a potential target for antifungal drugs to treat aspergillosis (46). Thus, RB-TDNAseq offers unparalleled potential for enriching genome annotation and elucidating gene function and interactions in filamentous fungi, improving our understanding of these ecologically, medically, and biotechnologically important organisms.

### Materials and Methods Strains and media

*Trichoderma atroviride* IMI 206040 was used as the wild-type strain for the generation of RB-TDNAseq libraries. *Agrobacterium tumefaciens* EHA105 was employed for barcoded library construction. For growth of the mutant library, we used both Potato Dextrose Broth (PDB) and Potato Dextrose Agar (PDA) media (BD Difco™).

### Reagents and filtration supplies

Acetosyringone (CAS No. 2478-38-8), hygromycin (CAS No. 31282-04-9), kanamycin (CAS No. 25389-94-0), and carbenicillin disodium salt (CAS No. 4800-94-6) were purchased from Millipore Sigma. For *Agrobacterium*-mediated transformation, we used HAWP MF-Millipore membrane filters (0.45 µm pore size; HAWP04700). All filtration steps were performed with Thermo Scientific™ Nalgene™ Rapid-Flow™ sterile disposable bottle-top filters equipped with polyethersulfone (PES) membranes. All primers used in this study were purchased from IDT, and their sequences are listed in Table S1.

### Randomly barcoded plasmid library construction

Generation of a large pool of uniquely barcoded plasmids was performed using an optimized type II-S endonuclease (Golden Gate) cloning strategy (24) modified from Coradetti *et al*. (14). The drug resistance cassette in the modified pGI2 (21) ATMT vector used in Coradetti *et al*. (14) (pDP11) was replaced with a hygromycin resistance cassette that expresses well in Sordariomycete species (Fig. 1). A *Hin*dIII recognition site in the region between the two divergent *Sap*I/*Lgu*I recognition sites was added just inside the right border of the TDNA in the pDP11 vector. We used these two divergent *Sap*I/*Lgu*I recognition sites to introduce the random barcodes. A population of random barcodes was generated by synthesizing the oligonucleotide GATGTCCACGAGGTCTCTNNNNNNNNNNNNNNNNNNNNCGTACGCTGCAGGTCGAC and amplifying it using the primers TCACACAAGTTTGTACAAAAAAGCAGGCTGGAGCTCGGCTCTTCGCCCGATGTCCACGAGGT CTCT and CTCAACCACTTTGTACAAGAAAGCTGGGTGGATCCGCTCTTCAATTGTCGACCTGCAGCGTACG (14). We combined 4 µg of plasmid DNA with 140 ng of barcode PCR fragments in a 50 µl reaction with 5 µl 10x T4 ligase buffer (ThermoFisher), 2.5 µl T4 ligase (ThermoFisher), and 2.5 µl *Lgu*I (ThermoFisher). We incubated the reaction for 5 m at 37°C, then 25 cycles of 2 m at 37°C, followed by 5 m at 16°C. The enzymes were heat inactivated for 10 m at 65°C. We let the product cool to 37°C, added 2.5 µl FastDigest Buffer (ThermoFisher), 2.5 µl Tango Buffer (ThermoFisher), 3.13 µl 2mg/ml bovine serum albumin (NEB), 12.5 µl FastDigest *Hin*dIII (ThermoFisher), and 12.5 µl *Lgu*I (ThermoFisher), and incubated the reaction at 37°C for 5 h to digest any uncut vector, then cooled to 4°C. The reaction products were purified using the Zymo DNA clean and concentrator-5 kit (Zymo Research) and eluted into 15 µl of ddH_2_O. To ensure a large diversity in the plasmid library, we pooled the products of 18 barcoded plasmid cloning reactions. We then transformed 25 µl of NEB 10-beta electrocompetent *E. coli* (NEB) with 5 µl of the Golden Gate cloning reaction product according to the manufacturer’s instructions in 48 independent transformations and selected for transformed *E. coli* using kanamycin. We pooled all transformants and estimated the diversity of the barcoded vector pool by sequencing plasmid barcodes on an Illumina HiSeq 4000 as described in Coradetti *et. al* (14), except the primers we used included dual indexes to prevent “index swapping” on the Illumina HiSeq 4000 (47) (48). The true pool size was estimated by the relative proportion of barcodes with 1 or 2 counts as described in the script Multicodes.pl (12). We estimate our barcoded plasmid pool contains approximately 1 billion uniquely barcoded plasmids (Dataset S1).

### *Agrobacterium tumefaciens* transformation with barcoded plasmid library

To generate a suitably diverse library of plasmids for *A. tumefaciens* mediated transformation of *T. atroviride*, we transformed the pooled barcoded plasmids into *A. tumefaciens* EHA105 using a method modified from (49). To generate electrocompetent *A. tumefaciens*, we inoculated *A. tumefaciens* into 5 ml LB and incubated at 30°C with shaking at 250 rpm for 16 h. We then back diluted the *A. tumefaciens* culture into 5 ml LB and incubated at 30°C with shaking at 250 rpm for at least 2 doublings until the cultures were at an optical density 600 (OD₆₀₀) of 1. We then back diluted a second time into 100 ml LB flasks such that they would be at OD₆₀₀ of 1 after an overnight incubation at 30°C with shaking at 250 rpm. *A. tumefaciens* cultures at OD₆₀₀ of 1 were chilled on ice with slow swirling for 45 m. These cultures were then decanted into ice-cold 50 ml conical tubes and spun at 4000 rpm for 15 m at 4°C. The cell pellet was then washed once in 50 ml ice-cold ddH_2_O, once in 25 ml ice-cold ddH_2_O, and once in 2 ml ice-cold 10% glycerol before resuspension in 150 µl of ice-cold 10% glycerol. Aliquots of 55 µl of cell suspension were then flash frozen in liquid nitrogen.

We transformed 153 aliquots of frozen cell suspensions. We thawed the frozen electrocompetent *A. tumefaciens* cells on ice and mixed the 55 µl cell suspension with 2 µg of barcoded plasmid DNA in ddH_2_O. We then incubated the cell/DNA mixtures on ice for 2 m before transferring the cell/DNA mixtures to chilled 2 mm gap 96-well electroporation plates (BTX) and electroporating using a 12.5 kV/cm field strength exponential decay wave (electroporation settings: 2500 V, 25 µF, 400 ohms) on a Gemini X2 high throughput electroporation system (BTX). We then immediately transferred the electroporated cells to 1 ml of 30°C LB and gently suspended the cells prior to incubating the culture for 2 h at 30°C with shaking at 250 rpm. Transformed *A. tumefaciens* cells were selected with kanamycin. We pooled all transformants, sequenced the barcodes, and estimated the diversity of the barcoded vector pool as described above. We estimate the barcoded plasmid pool housed in *A. tumefaciens* contains over 140 million uniquely barcoded plasmids (Dataset S1).

### A. tumefaciens mediated transformation of T. atroviride

We grew the barcoded *A. tumefaciens* pool to an OD₆₀₀ of 1 in 50 mL of LB with kanamycin in a baffled flask at 30°C at 250 rpm overnight. We then centrifuged the cells, resuspended them in 10 mL of induction medium (ABI medium - described below), and incubated them for 24 h at room temperature in culture tubes on a roller drum. To generate *T. atroviride* conidia, we cultured *T. atroviride* on PDA for 84 h in constant light. After harvesting the conidia with sterile water, 2 × 10^7^ *T. atroviride* conidia were pelleted by centrifugation at 4,000 rpm for 15 min, and the supernatant was discarded. The pellet was then resuspended in 2 ml of the induced *A. tumefaciens* culture at an OD₆₀₀ of 1 and incubated for 5 min at room temperature. The *T. atroviride*-*A. tumefaciens* mixed culture was then filtered onto a 0.45 µm pore membrane filter, with a diameter of 47 mm. For our library construction, we used a total of 100 filters to transform ∼ 2 x 10^9^ *T. atroviride* conidia. The filter with the cells was transferred onto petri plates containing ABI medium with 2% bacto agar with the cells facing up and incubated at 21°C for 7 days in darkness.

To induce *A. tumefaciens*-mediated transformation of *T. atroviride* conidia, we prepared a modified co-cultivation medium (ABI medium) based on AB minimal medium (50). The ABI medium is composed of several stock solutions: 20x AB salts solution, 500 mM phosphate buffer at pH 7.5, a 2x 2-(N-morpholino)ethanesulfonic acid (MES) buffer at pH 5.0, and a 500 mM acetosyringone solution. The 20x AB salts contained 20 g/L ammonium chloride (NH₄Cl), 6 g/L magnesium sulfate heptahydrate (MgSO₄·7H₂O), 3 g/L potassium chloride (KCl), 0.2 g/L calcium chloride (CaCl₂), and 15 mg/L iron (II) sulfate heptahydrate (FeSO₄·7H₂O) and was filter sterilized. The 500 mM phosphate buffer contained 60 g/L dipotassium phosphate (K₂HPO₄) and 20 g/L sodium dihydrogen phosphate (NaH₂PO₄), with pH adjusted to 7.5 using 2 M KOH. The 2× MES buffer contained 9.72 g/L MES, with pH adjusted to 5.0 with 2 M KOH. The 500 mM acetosyringone solution contained 0.098 g/mL of acetosyringone in dimethyl sulfoxide (DMSO) and was either freshly prepared or aliquoted and stored at –20 °C. The stock solutions were mixed with 50 mL/L 20x AB salts solution, 2.4 mL/L 500 mM phosphate buffer, 500 mL/L 2x MES buffer, and 1 mL/L 500 mM acetosyringone (added immediately prior to use). This mixture was supplemented with 0.2% glucose (w/v), 0.05% casamino acids (w/v), and 0.001% thiamine (w/v), and the solution was stored at 4 °C. All components were sterilized separately, then mixed under sterile conditions, and either used immediately or stored at 4°C until needed.

After a 7-day co-cultivation of *T. atroviride* conidia with *A. tumefaciens* on filters on ABI solid medium, we harvested the transformed *T. atroviride* conidia by vigorously vortexing the 0.45 μm membrane filter in 5 mL of sterile ddH_2_O to detach the cells. We transferred 2 mL of the suspension to a sterile 2 mL Eppendorf tube and centrifuged at 10,000 rpm for 5 minutes. The supernatant was carefully discarded, and the remaining suspension was added to the same tube and subjected to a second round of centrifugation under the same conditions. After discarding the final supernatant, the resulting pellet was resuspended in 1 mL sterile ddH_2_O.

To select for transformed *T. atroviride* cells without contamination by untransformed *T. atroviride* cells or *A. tumefaciens* cells, we used a selection system consisting of multiple layers of agar in 150 mm petri dish (Fig. S3). In total we used ten 150 mm petri dishes. We first poured a layer (Layer 1) of 50 mL PDA + 200 µg/mL hygromycin (to select for transformed *T. atroviride*) + 100 µg/mL carbenicillin (to select against *A. tumefaciens*). A 13.5 ml PDA + 100 µg/mL carbenicillin (Layer 2) was then poured over the first solidified layer. Once the second layer solidified, we then spread 10 mL of the conidial suspension (harvested from the filter as described above) on top of Layer 2 (Layer 3). After the conidial suspension dried, we poured 13.5 mL of 37°C low-melting-point PDA+ with 100 µg/mL carbenicillin, over the conidial suspension (Layer 4). Once layer 4 solidified, we poured 13.5 mL of 55°C PDA+ with 200 µg/mL hygromycin + 100 µg/mL carbenicillin as the final layer (Layer 5) to preserve selective pressure on *T. atroviride* (Fig. S3). Inoculated plates were incubated at 28°C for 6 days in constant light to induce conidiation. Only fungal colonies capable of growing invasively through Layers 4 and 5 were recovered for downstream analyses.

Transformed *T. atroviride* conidia were harvested and resuspended in 10 mL of ddH_2_O. The suspensions were pooled and 50 mL aliquots were transferred to 50 mL Falcon tubes and centrifuged at 4,100 rpm for 15 m to concentrate the conidia. The pellet was then resuspended in 10 mL of sterile 30% (v/v) glycerol per each Falcon tube and stored at –80°C for long-term preservation. We collected transformants from ten 150 mm Petri dishes, corresponding to the output of 100 filters on which the transformation was performed (10 filters of transformants per plate). The protocol is available at: https://www.protocols.io/view/transformation-protocol-atmt-trichoderma-atrovirid-261gekk87g47/v1.

### Genomic DNA extraction from the TDNA library of *T. atroviride*

Genomic DNA was extracted from the *T. atroviride* TDNA library conidia using the Quick-DNA™ Fungal/Bacterial Miniprep Kit (Zymo Research, catalog no. D6005). Conidia were collected in sterile ddH_2_O (200 μL of liquid with 10^9^ conidia) and resuspended in 500 µL of Genomic Lysis Buffer. The suspension was transferred to a ZR BashingBead™ Lysis Tube, followed by the addition of 750 µL of BashingBead™ Buffer. The tubes were placed on ice and then processed in a bead beater at maximum speed (Mini-Beadbeater-96) for 2 min. The remaining steps were carried out following the manufacturer’s protocol. To assess DNA integrity, we ran the genomic DNA on a 1% agarose gel, looking for a single band without schmearing. To assess DNA purity, we quantified the DNA using both a NanoDrop spectrophotometer and a Qubit fluorometer to confirm that the DNA did not have RNA contamination.

To quantify the reduction of residual *A. tumefaciens* DNA, we performed a qPCR-based assay on genomic DNA from transformed *T. atroviride* conidia (Fig. S3C). qPCR was conducted on a Bio-Rad CFX96 real-time system (96-well format). *A. tumefaciens* DNA was quantified using primers specific to the *aopB*gene (Forward: CTGGTTAGGCTTCTGTTAGGG; Reverse: GGAGAATGCGGAGATCAAAGAG), which amplify a 118-bp fragment. For normalization, we amplified the *T. atroviride* actin gene (Gene ID: 297070) with primers Forward ACCTCTACGGCAACATTGTC and Reverse CAGTGATCTCCTTCTGCATACG, yielding a 140-bp product.

### TDNAseq library preparation

To link barcode sequences to their specific genomic locations within the conidia pool, we sequenced the sites of TDNA insertions (TDNAseq) by adapting a modified version of the KAPA DNA sequencing library preparation protocol (29). Briefly, 1 µg of high-quality genomic DNA was sheared to an average size of 300 bp using a Covaris sonicator compatible with microTUBE AFA Fiber Crimp-Cap tubes (6×16 mm; Covaris, cat. no. 520052 or 520091), in the presence of pH 8 TE buffer. Fragmented DNA was size-selected through a double-sided solid-phase reversible immobilization (SPRI) procedure using AMPure XP magnetic beads (Beckman Coulter, cat. no. A63881) and freshly prepared 80% ethanol for wash steps. The resulting DNA fragments underwent end-repair and A-tailing using the KAPA Hyper Prep Kit (Roche, cat. no. KK8502) according to the manufacturer’s instructions. Adapter ligation was performed in the same reaction tubes using pre-annealed splinkerette adapters and the ligation reagents included in the KAPA kit. The ligation products were purified using a one-sided SPRI bead cleanup.

Amplification of the adapter-ligated DNA was carried out in two successive PCR steps using KAPA HiFi HotStart ReadyMix (Roche, cat. no. KK2601). The first PCR selectively amplified fragments containing TDNA-genome junctions, and the product was cleaned using a double-sided SPRI selection to remove short and nonspecific products. A second PCR was then performed to incorporate unique Illumina-compatible index sequences into each sample and select for products containing a TDNA insertion, followed by another double-sided SPRI cleanup to eliminate adapter dimers and undesired fragment sizes. The final library quality and fragment size distribution were assessed using an Agilent 2100 Bioanalyzer with the DNA 1000 Kit (Agilent Technologies, cat. no. 5067-1504). Only libraries displaying a clear peak around 400 bp and minimal adapter contamination were retained for high-throughput sequencing. Sequencing was performed using 150 bp paired-end reads across two lanes on an Illumina MiSeq platform at QB3 Genomics, UC Berkeley.

## Mapping insertion locations

The location of barcoded TDNA insertions in the *T. atroviride* RB-TDNA library were mapped to the *T. atroviride* genome using the GFF annotations available at the NCBI (https://www.ncbi.nlm.nih.gov/datasets/taxonomy/63577/), as described (14, 51) (https://github.com/stcoradetti/RBseq). The specific settings for the analysis with *T. atroviride* are in https://github.com/jmvillalobos/Trichoderma_RB-TDNAseq. To filter true barcodes present in our library from sequencing errors, we eliminated any barcodes that differed from another, much more abundant, barcode by only one or two bases. We also eliminated barcodes whose insertion mapped to more than one location in the *T. atroviride* genome, suggesting the respective TDNA had been inserted into multiple locations in one or many transformants.

## GO term enrichment analysis

We used gene annotation reported for *T. atroviride* (30, 52). To perform the functional category enrichment analysis, we used Gene Ontology (GO) terms and conducted the analysis with the GOseq 1.61.1 package in R (53) (Dataset S3). Enrichment and depletion were assessed using the set of all annotated genes in the genome as the background universe, and as the test set, we considered genes containing at least one insertion.

## Data and Code availability

Sequencing data are available on the National Center for Biotechnology Information (NCBI) Sequence Read Archive (SRA) with accession number PRJNA1347553 (https://www.ncbi.nlm.nih.gov/bioproject/PRJNA1347553). All custom scripts and code used in the analysis are available at: https://github.com/jmvillalobos/Trichoderma_RB-TDNAseq and https://github.com/stcoradetti/RBseq.

## Acknowledgements

This material by m-CAFEs Microbial Community Analysis & Functional Evaluation in Soils, a Science Focus Area led by Lawrence Berkeley National Laboratory, is based upon work supported by the U.S. Department of Energy, Office of Science, and Office of Biological & Environmental Research under contract number DE-AC02-05CH11231. This work was also supported by a University of California Berkeley Innovative Genomics Institute grant to N.L.G., A.P.A., and J.M.S., a grant from the National Institute of General Medical Sciences of the National Institutes of Health under award number R35GM150926 to L.B.H, and support from the Challenge-Based Research Funding Program under award number IJXT070-23EG54001 to J.M.V.E. We thank Alfredo Herrera-Estrella (Center for Research and Advanced Studies, Irapuato, Guanajuato, Mexico) for his help in designing the conidia experiments. The authors also wish to thank María Belén Mercado-Esquivias for her support in writing the transformation protocol and for assistance with the ATMT experiments and Ya-Fang Cheng for her assistance with optimizing ATMT of filamentous fungi.

