## Supplemental Figures and Tables for "Construction of a randomly barcoded insertional mutant library in the filamentous fungus *Trichoderma atroviride*"

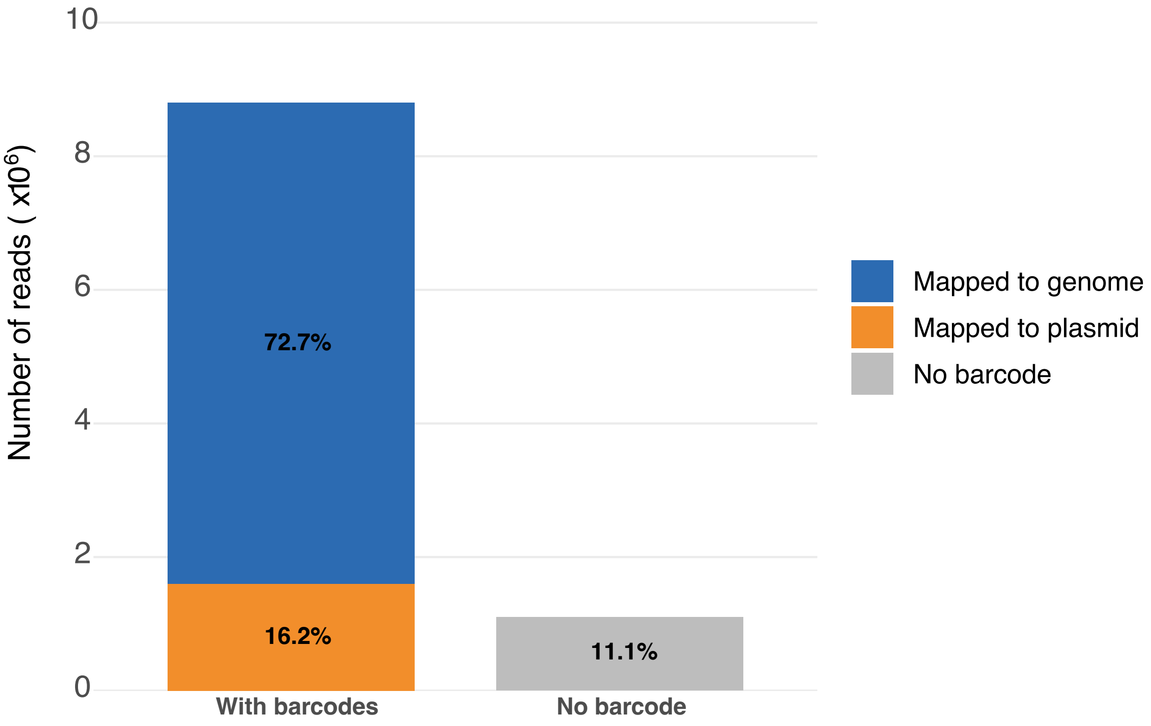


**Figure S1. Distribution of reads with and without a barcode.** Approximately 72.7% of reads map to insertions in the *Trichoderma atroviride* genome. Approximately 16.2% of reads map to the plasmid (indicative of bacterial contamination, incorrect processing of the plasmid during ATMT, or concatemer formation due to back-to-back insertion of TDNA), and only 11.1% of reads lack barcodes.

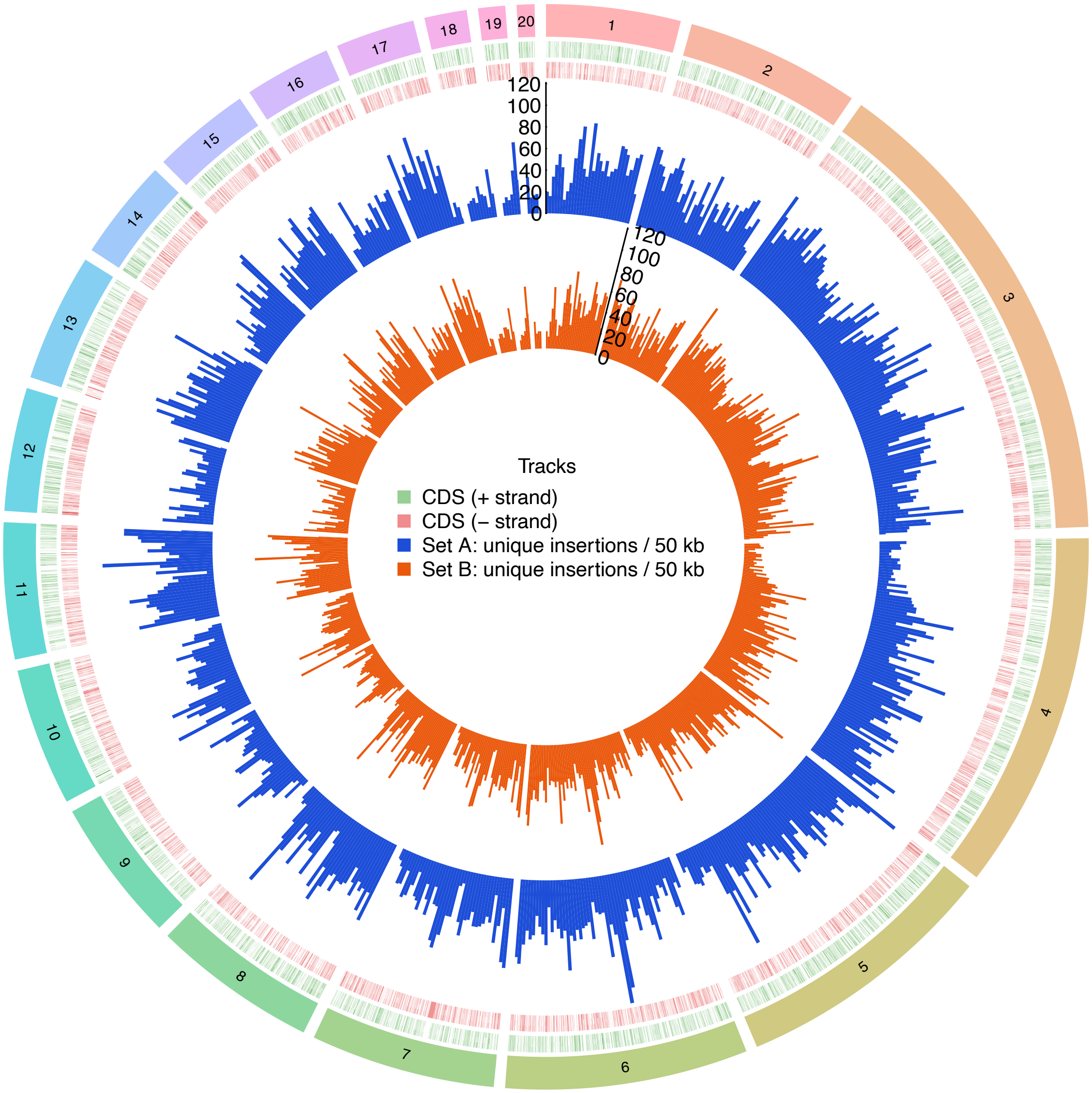


**Figure S2. Circos plot showing the density of unique barcode insertions identified in two TDNAseq sequencing runs from the same pool of transformants.** The KAPA library prep was performed separately on the same insertional mutant library for the two sequencing runs (Set A, blue; Set B, orange). Each bar represents the number of unique TDNA insertion sites within non-overlapping 50 kb genomic windows. Contigs are numbered sequentially (1–20) according to their genomic size. The y-axis indicates the absolute number of unique insertions per 50 kb window, and both libraries share the same scale for direct visual comparison. Gene coding sequences (CDS) are shown in green (positive strand) and red (negative strand). This visualization highlights regional differences in TDNA insertion density, revealing hot and cold spots of TDNA integration. The plot was generated using the R package circlize (v0.4.16) (1).

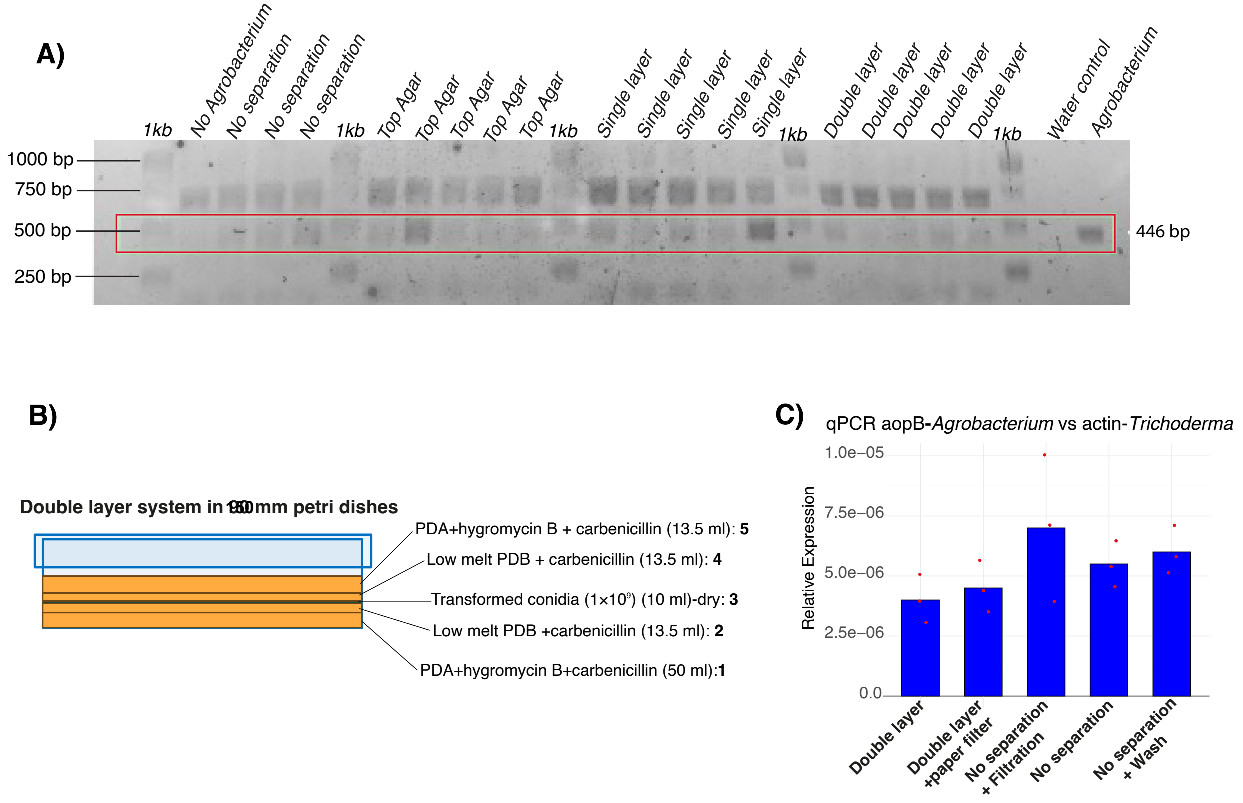


**Figure S3: Verification of the reduction of *Agrobacterium tumefaciens* DNA during the recovery of the transformed conidial library.** (**A**) PCR amplification of the *aopB* (outer membrane protein) gene, specific to bacteria and absent in fungi, which should result in a 446 bp PCR product. The DNA ladder used is 1 kbp Plus from Thermo Scientific (labeled as "1kb" on gel). (**No *Agrobacterium*:** Untransformed conidia. **No separation:** Transformed conidia were plated directly on PDA + hygromycin B + carbenicillin and conidia were recovered after incubation. **Top agar:** A layer of PDA + hygromycin B + carbenicillin was poured first. Then a layer of PDA + carbenicillin was poured over that. The transformed conidia were plated onto the PDA + carbenicillin layer and conidia were recovered after incubation. **Single layer:** After the conidia were inoculated onto the top agar, a layer of low-melt PDB agar + carbenicillin was added, and conidia were recovered after invasive growth through this final layer. **Double layer:** Single layer system with a second layer of PDA + hygromycin B + carbenicillin added on top of the top agar, and conidia were recovered after invasive growth through this uppermost layer (i.e., two additional layers above the top agar).) (**B**) Schematic representation of the layered approach used during library construction to control bacterial contamination in the *T. atroviride* library. (**C**) qPCR analysis of bacterial DNA concentration in the *T. atroviride* library, shown as the amount of the *A. tumefaciens aopB* gene relative to the amount of the actin gene from *T. atroviride*. Although differences were not statistically significant, the double-layer method had generally lower *aopB* to *actin* ratios than other methods. (**Double layer:** The system shown in (**B**). **Double layer + paper filter:** The system shown in (**B**) with a sheet of filter paper placed on top. The mycelia grew through the agar and the filter paper and conidia were collected from the filter paper. **No separation + Filtration:** Transformed conidia were plated directly on PDA + hygromycin B + carbenicillin and conidia were collected after incubation. The conidial suspension was then filtered through a Whatman filter and the conidia that were retained on the Whatman filter were recovered again by resuspending them in water. **No separation:** Transformed conidia were plated directly on PDA + hygromycin B + carbenicillin and conidia were collected after incubation. **No separation + Wash:** Transformed conidia were plated directly on PDA + hygromycin B + carbenicillin, conidia were collected after incubation, centrifuged once, the supernatant was discarded, and the conidia were resuspended in water.)

**Table S1.** Primers used in this study and their applications.

| **Gene/Purpose** | **Forward primer (5'->3')** | **Reverse primer (5'->3')** | **Notes** |
|---|---|---|---|
| *aopB* (*A. tumefaciens* qPCR marker) | CTGGTTAGGCTTCTGTTAGGG | GGAGAATGCGGAGATCAAAGAG | Used to detect bacterial contamination by qPCR |
| Actin (*T. atroviride* qPCR marker) | ACCTCTACGGCAACATTGTC | CAGTGATCTCCTTCTGCATACG | *T. atroviride* internal control for qPCR normalization |
| Amplification of random barcodes for Golden Gate cloning | TCACACAAGTTTGTACAAAAAAGCAGGCTGGAGCTCGGCTCTTCGCCCGATGTCCACGAGGTCTCT | CTCAACCACTTTGTACAAGAAAGCTGGGTGGATCCGCTCTTCAATTGTCGACCTGCAGCGTACG | Primers used to amplify fragments containing random barcodes for Golden Gate cloning (2) |
| Amplify TDNA-genome junctions for TDNAseq | AATGATACGGCGACCACCGAGATCTACACTCTTTCCCTACACGACGCTCTTCCGATCTNNNNNNcccgatgtccacgaggtctct | CAAGCAGAAGACGGCATACGAGATCGTGATGTGACTGGAGTTCAGACGTGTGCTCTTCCGATCT | Primers for second round PCR of the TDNAseq library prep |
| Amplify TDNA-genome junctions for TDNAseq | CTCCACTAGCTCCAGCCAAG | GAGATCGGTCTCGGCATTC*C | Primers for first round PCR of the TDNAseq library prep |

**Supplemental Datasets:**

**Dataset S1.** The number of barcodes sequenced in the **A**) *Escherichia coli* or **B**) *A. tumefaciens* plasmid libraries indicated as the number of barcodes with a given number of sequencing reads. The Chao estimate of the true library size is estimated by the relative proportion of barcodes with 1 or 2 counts as described in the script Multicodes.pl (3).

**Dataset S2.** Annotation of genomic regions with insertions (exon, intron, promoter, terminator, and intergenic) from two independently sequenced preparations of the same insertional mutagenesis library (**A** and **B**). The dataset reports the number of genomic nucleotides represented by each category and the number of insertions mapped to each category.

**Dataset S3.** Functional category analysis (4) of genes with at least one TDNA insertion in the *T. atroviride* barcoded insertional mutant library.
